# 3D Visualisation of trans-syncytial nanopores provides a pathway for paracellular diffusion across the human placental syncytiotrophoblast

**DOI:** 10.1101/2022.01.26.477815

**Authors:** Rohan M Lewis, Harikesan Baskaran, Jools Green, Stanimir Tashev, Eleni Paleologou, Emma M Lofthouse, Jane K Cleal, Anton Page, David S Chatelet, Patricia Goggin, Bram G Sengers

**Author notes:** **Corresponding Author:** Rohan M Lewis, MP 887, IDS Building University of Southampton, Faculty of Medicine, Southampton General Hospital.

## Abstract

The placental syncytiotrophoblast, a syncytium without cell-cell junctions, is the primary barrier between the mother and the fetus. Despite no apparent anatomical pathway for paracellular diffusion of solutes across the syncytiotrophoblast size-dependent paracellular diffusion is observed. Here we report data demonstrating that the syncytiotrophoblast is punctuated by trans-syncytial nanopores (TSNs). These membrane-bound TSNs directly connect the maternal and fetal facing sides of the syncytiotrophoblast, providing a pathway for paracellular diffusion between the mother and fetus. Mathematical modelling of TSN permeability based on their 3D geometry suggests that 10-37 million TSNs per cm^3^ of placental tissue could explain experimentally observed placental paracellular diffusion. TSNs may mediate physiological hydrostatic and osmotic pressure homeostasis between the maternal and fetal circulations but also expose the fetus to pharmaceuticals, environmental pollutants and nanoparticles.

## Introduction

The placenta was once viewed as a perfect barrier, but as the thalidomide tragedy demonstrated, this is not the case [1]. It is now clear that potentially harmful molecules and particulates can cross the placenta and adversely affect fetal development. However, the mechanism by which these molecules and particulates cross the placenta is not always clear [2, 3]. Understanding how these substances cross the placenta is necessary to identify the risks and prevent long-term consequences of these exposures, which may adversely affect fetal and postnatal health [4].

The primary placental barrier is the syncytiotrophoblast, a continuous syncytial monolayer covering the villi at the maternal-fetal interface. As there are no cell-cell junctions in a syncytium, there is no obvious pathway by which paracellular diffusion can occur. Despite the absence of an anatomical pathway for diffusion, there is physiological evidence for size-dependent paracellular diffusion of solutes [5, 6]. Trans-syncytial channels, or nanopores, have been proposed as mediators of trans-syncytial diffusion, however continuous full-width nanopores have not been previously demonstrated in the human placenta [7]. An alternative hypothesis to explain paracellular diffusion is that it occurs through regions of syncytial damage [8]. These hypotheses are not mutually exclusive but establishing mechanisms of fetal exposure is necessary to understand the likely risks of different compounds and develop strategies to mitigate this.

Selective placental transfer of nutrients, IgG, wastes and exogenous toxins is facilitated by membrane transporters and endocytosis [9-11]. However it is not clear how exogenous drugs and toxins reach the fetus as, with a few exceptions such as the apically located exchanger OATP4A1 [10], drug transporters in the placenta mediate efflux from the fetus to the mother [12]. Nanoparticle transfer across the placenta has been observed, but the mechanism is unclear [13]. A more extensive understanding of how metabolites, pharmaceuticals and toxins reach the fetal circulation is necessary to protect fetal health.

Estimates of placental permeability surface area products for hydrophilic solutes have been determined *in vivo* and show size-dependent permeability of solutes, which decreases with increasing molecular radius [5, 6]. In other species with haemochorial placentas, permeability has also been shown to be size-selective with some species having higher or lower overall permeability compared to humans [3, 14]. This data suggests a size-selective permeability of the placenta through low diameter channels [15].

Using serial block-face scanning electron microscopy (SBF SEM) to reconstruct placental ultrastructure in three dimensions, this study demonstrates the presence of full-width trans-syncytial nanopores (TSNs) in the human placenta.

## Methods

Term placental tissue was collected after delivery from uncomplicated pregnancies with written informed consent and ethical approval from the Southampton and Southwest Hampshire Local Ethics Committee (11/SC/0529). Villous samples from 8 placentas were processed and imaged using TEM or SBFSEM as described previously [27]. Regions containing TSNs were manually segmented in Avizo v2019.4 (ThermoFisher, Eindhoven).

### Modelling of nanopore diffusive transfer capacity

The nanopore diffusive transfer capacity for a particular molecule is determined by its diffusivity in free solution as well as the geometry of the pore, specifically the pore length and cross sectional area. Here we will calculate this geometric contribution in the form of the effective area-over-length ratio *A_pore_/L_pore_* [m], which is a measure for the effect of pore geometry on diffusive transfer, independent of the solute studied. This parameter can then be used to calculate nanopore diffusive transfer using the concentration gradient and diffusion coefficient for any specific solute.

Briefly, the magnitude of the diffusive flux *J_pore_* [mol/s] through a pore of uniform cross section is given by:

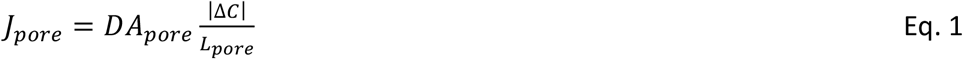

where *D* [m^2^/s] is the diffusion coefficient, *A_pore_* [m^2^] the pore cross sectional area, Δ*C* [mol/m^3^] the concentration difference over the pore and *L_pore_* [m] the pore length. This can also be expressed in terms of the permeability surface area product *PS_pore_* [m^3^/s]:

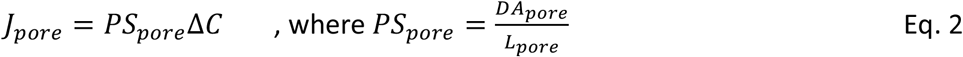

Since in reality the pore cross sectional area is not uniform, 3D image-based modelling was used to calculate the effective area-over-length ratio *A_pore_/L_pore_* [m] based on Eq. 1 by prescribing the concentration difference and diffusivity and then monitoring the resulting flux in the simulations. Segmented image stacks were imported in Simpleware ScanIP (P-2019.09; Synopsys, Inc., Mountain View, USA). Voxels were resampled isotropically so that the x and y pixel size matched the z axis spacing. Pores were meshed using linear tetrahedral elements. Different coarseness settings were evaluated to establish convergence, with the final number of elements used for the different pores ranging from 1.4-38×10^5^.

Steady state diffusion simulations were performed in COMSOL Multiphysics (v5.5; COMSOL AB, Stockholm, Sweden). Since *A_pore_/L_pore_* does not depend on the choice of parameters, *D* = 1 m^2^/s was used and a fixed concentration gradient ΔC = 1 mol/m^3^ was imposed by constant concentration boundary conditions on the inlet and outlet surface of the pore, while the remaining external surface was subject to no-flux conditions.

After simulations, the magnitude of the diffusive flux *J_pore_* given by the integral of the solute flux in the direction normal to the pore inlet or outlet taken over the inlet/outlet cross sectional area, was then used to calculate the effective pore area-over-length ratio *A_pore_/L_pore_* based on Eq. 1.

For a specific solute, the permeability surface area product for a single pore *PS_pore_* was obtained by multiplying the average *A_pore_/L_pore_* value by the relevant diffusion coefficient (Eq. 2). The number of pores *n_pores_* [g^−1^] per gram of placental tissue was then estimated based on Eq. 3:

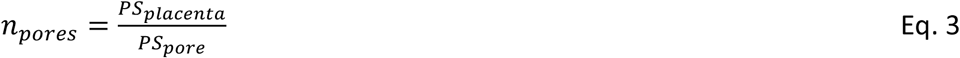

where *PS_placenta_* [m^3^ s^−1^ g^−1^] is the overall experimentally observed placental permeability surface area product for the corresponding solute from literature.

## Results

### Trans-syncytial nanopores

This study manually inspected 14 SBFSEM image stacks from five different placentas to identify trans-syncytial nanopores (TSNs) defined as membrane lined pores connecting the apical and basal plasma membranes of the syncytiotrophoblast. These stacks consisted of 7487 SBFSEM images, representing a total volume of 0.00000017 cm^3^.

Inspection of serial sections allowed identification of TSNs crossing the syncytiotrophoblast. Continuous TSNs with a clear connection between the apical and basal membranes were identified. Near-continuous TSNs were also identified which had connections to both the apical microvillous and basal plasma membranes but contained discontinuities along the length of the nanopore. Unilateral nanopores, ultrastructurally similar to TSNs but opened from either the apical or basal plasma membrane without connecting to the opposite membrane were identified. An illustration of these nanopore types as we defined then is shown in (figure 1). Regions containing TSNs were manually segmented in Avizo v2019.4 (ThermoFisher, Eindhoven). Cross section from one of these TSNs are shown in figure 2. The lumens of TSNs typically had low electron density, consistent with a primarily fluid filled pore (figure 2a, c and d). However, some TSNs and regions of TSNs were observed with higher electron density indicating diffuse contents (figure 2e). The nanopore shown in figure 2 can be seen reconstructed in 3D (Supplemental movie 1) and a movie showing the nanopore lumen highlighted in sequential sections (supplemental movie 2) can be seen in the supplemental material.

**Figure 1.**
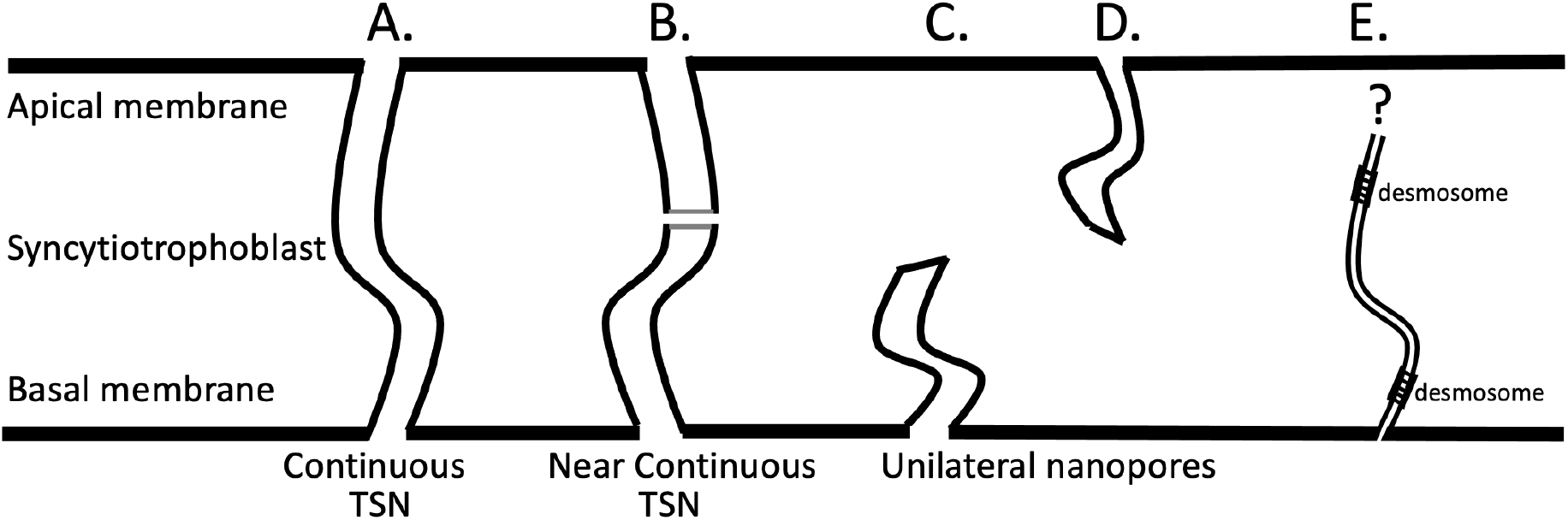
Classification of nanopores found in human placental syncytiotrophoblast. **A)** continuous TSNs connecting the apical and basal membrane without any breaks. **B)** near-continuous TSNs with sections connected to the apical and basal plasma membranes but contained discontinuities where no clear connection could be observed but the ends of the discontinuous sections were adjacent. **C & D)** Unilateral nanopores were ultrastructurally similar to TSNs but opened from either the apical or basal plasma membrane but without any apparent connection to the opposing membrane. **E)** Desmosome associated nanopores were also observed arising from the basal membrane but none of these could be observed connecting to the apical membrane.

**Figure 2.**
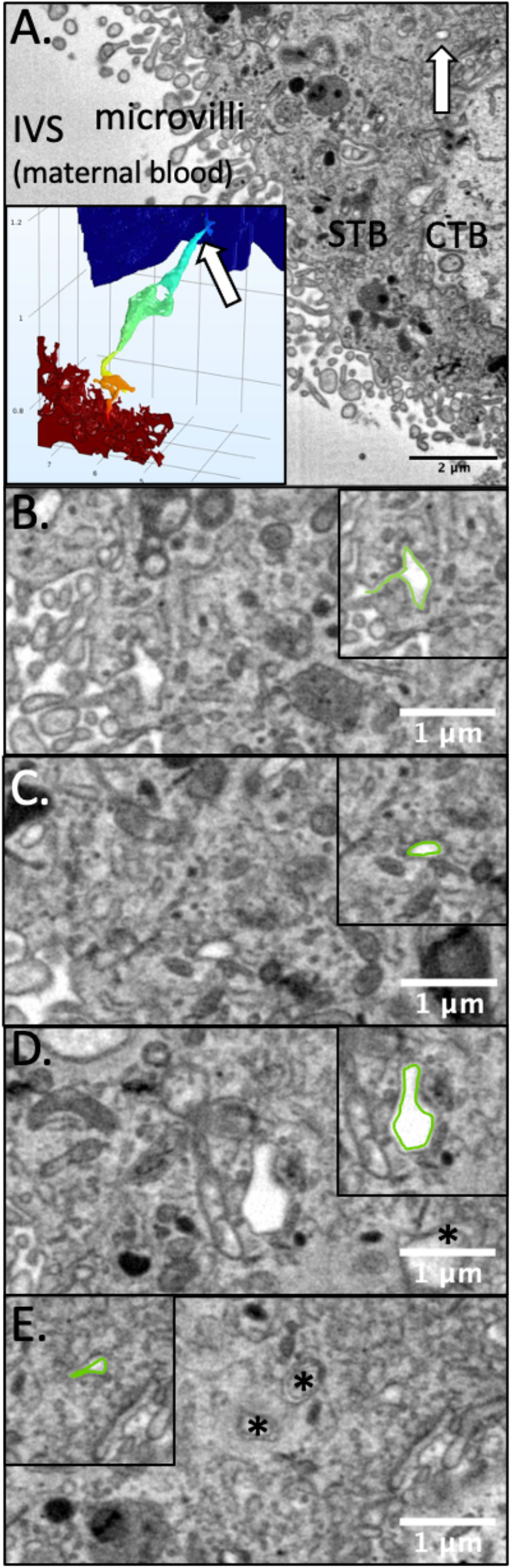
SBF SEM serial sectioning allows identification of TSNs crossing the human placental syncytiotrophoblast. **A)** An SBF SEM image showing a cross section of the syncytiotrophoblast (STB) from the maternal intervillous space (IVS) to the underlying cytotrophoblast (CTB). Within this image a cross section of a nanopore can be seen (white arrow) illustrating how difficult it would be to identify these structures from individual 2D images. The inset shows the nanopore reconstructed in 3D from 44 consecutive sections with the white arrow indicating where the section comes from and modelled solute concentration indicated by colour from high (red) to low (blue). **B)** the apical opening of the TSN with the inset showing this highlighted in green. **C)** A thin region of nanopore with the inset showing this highlighted in green. **D)** a dilated region of the nanopore with the inset showing this highlighted in green. In B, C and D the nanopore has a low electron density consistent with a fluid filled pore **E)** A region of nanopore where the lumen has a higher electron density than in other regions. The inset showing this cross section of the nanopore highlighted in green. This region is close to the end of the nanopore and the cytotrophoblast boundary can be seen bottom left. * Indicates inclusions in other TSNs in the section. A movie showing the nanopore reconstructed in 3D and another movie with the nanopore lumen highlighted in sequential sections can be seen in the supplemental material.

Ten continuous TSNs were identified from three different placentas and these did not have a uniform structure (figure 3). While the least complex TSNs were direct tubes connecting the apical and basal membranes (figure 3c) the most complex full width TSN was a double pore with two bifurcated apical openings (4 in total) and two basal openings (figure 3a). Blind ends are present in the TSNs shown in figure 2d, i and g.

**Figure 3.**
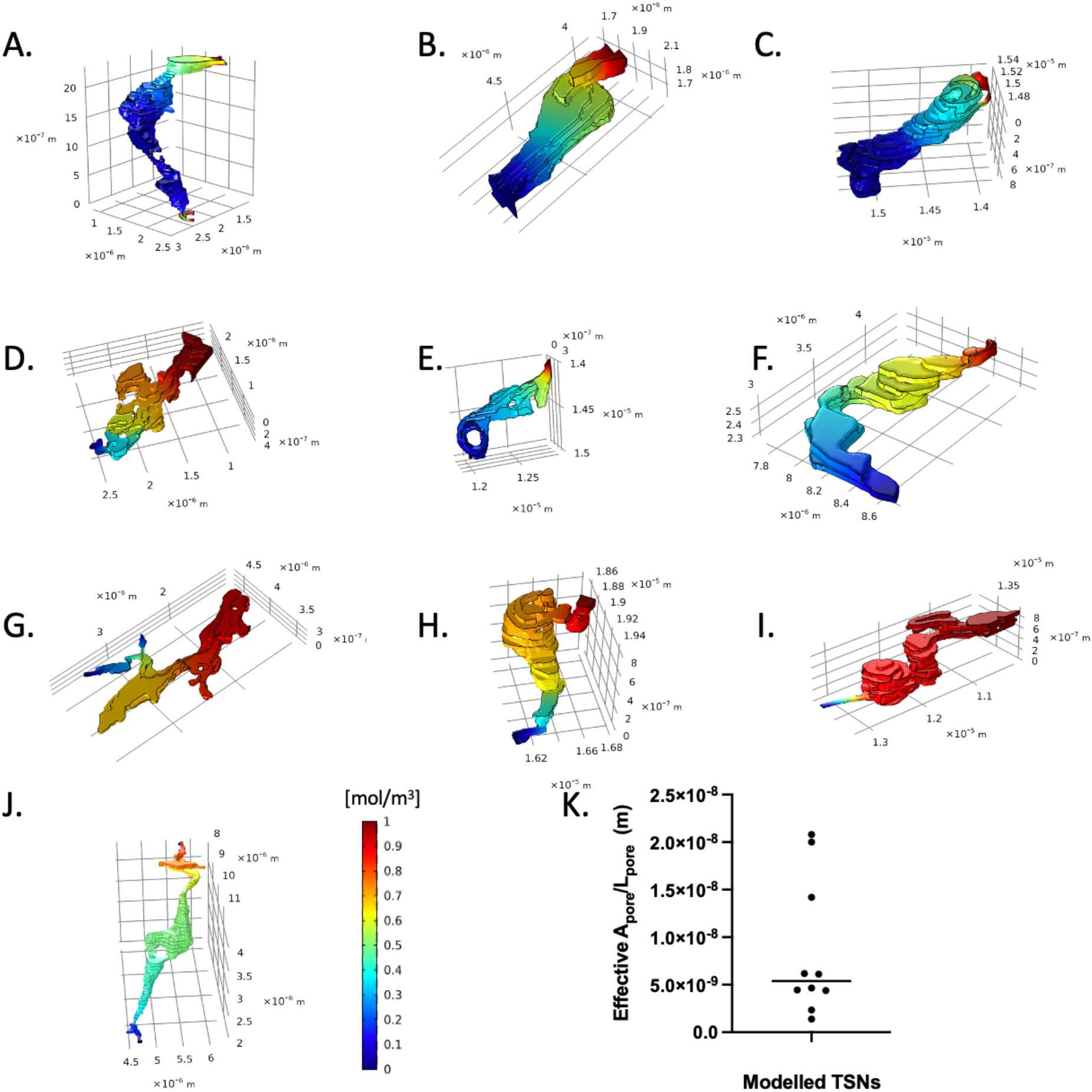
Modelled solute concentration gradients within the continuous TSNs. High concentrations (red) were applied to the maternal facing TSN opening, and concentration gradients are shown for diffusion through the channel. The molecular flux associated with these gradients was used to calculate the effective cross sectional surface area used to calculate permeability. **A-J)** show the nanopores presented in order of calculated permeability from highest to lowest. The apical opening is presented on top right and basal openings on the bottom or left. The TSN shown in A is a double pore with four apical and basal openings and a connection on the basal side which accounts for its higher permeability despite its longer length. Blind ends are present in D, I and G. TSN A has two openings to the microvillous membrane and two openings to the basal membrane of the syncytiotrophoblast. **J)** shows a scatterplot showing the distribution of permeabilities with the line indicating the median value.

In addition, 20 near-continuous TSNs were identified, with examples in all five placentas studied. In two cases, these near-continuous TSNs were highly complicated with branches, multiple dilations, and blind ends (figure 4b). Finally, 25 unilateral nanopores were identified arising from either the apical (n = 17) or basal (n = 8) plasma membrane of the syncytiotrophoblast but not connecting to the opposing plasma membrane.

**Figure 4.**
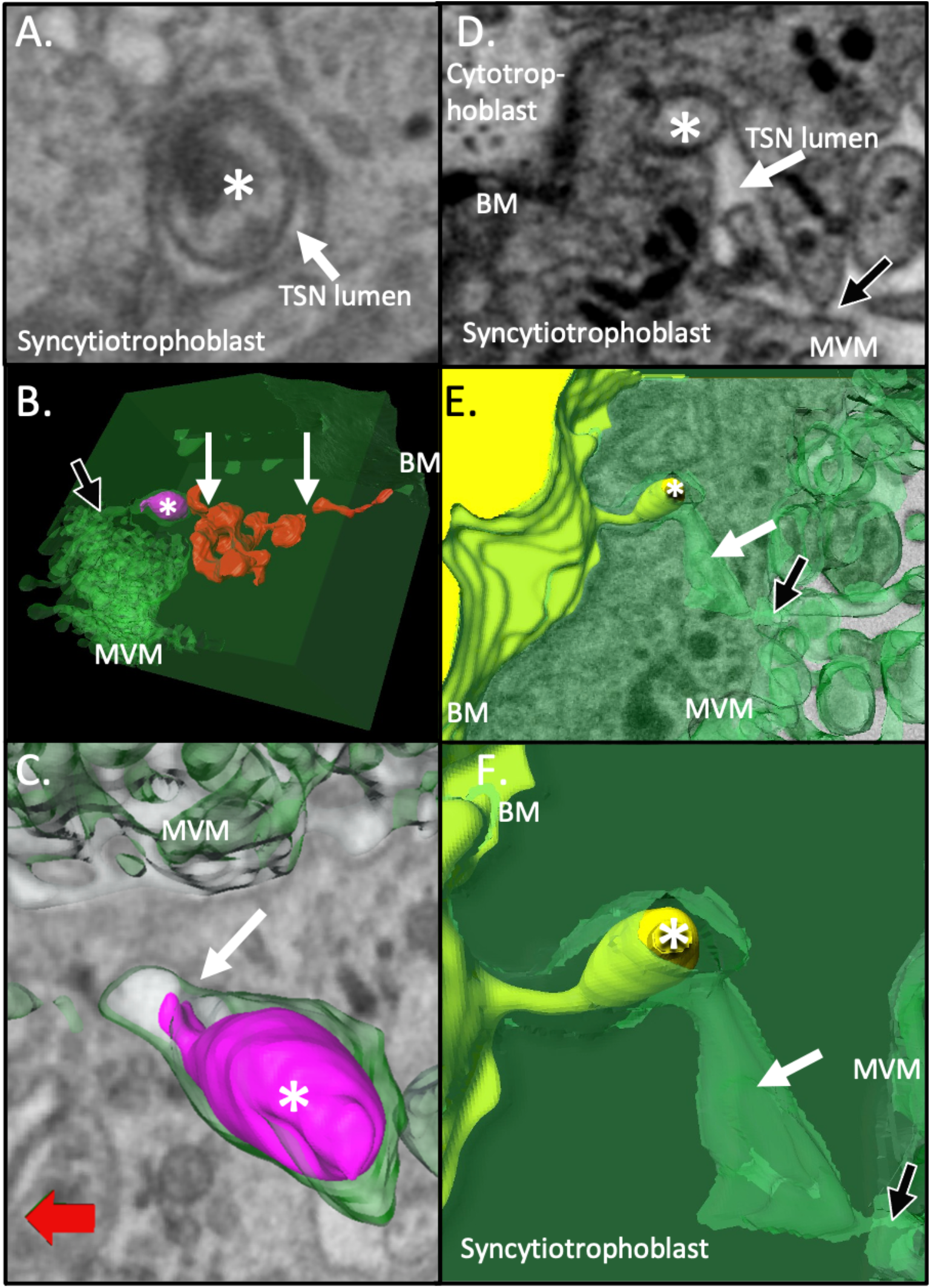
Examples of TSN inclusions formed from engulfed syncytiotrophoblast or villous cytotrophoblast. **A-C**) An example of an apical nanopore inclusion (highlighted by an *) which in 3D can be seen to be budding off the syncytiotrophoblast within the nanopore. **A)** shows a SBF SEM image containing a cross section of the TSN inclusion. **B)** shows the pink inclusion reconstructed in 3D within the TSN formed from green syncytiotrophoblast. This inclusion lies within a complex near-continuous TSN which, apart from regions around the inclusion, is shown in red. This TSN has discontinuities indicated by white arrows but gives the appearance of being part of a trans-syncytial network. A black arrow indicates the apical opening of this TSN. **C)** shows a higher magnification image of the pink inclusion and the connection to the green syncytiotrophoblast is indicted by a white arrow. **D-F)** An example of a TSN inclusion, which in 3D is seen to be derived from underlying villous cytotrophoblast. In panels D-F an * indicates the inclusion, white arrows indicate the lumen of the TSN and black arrow indicates the apical opening of the TSN on the microvillous membrane. **D)** shows a 2D SBF SEM image of the syncytiotrophoblast containing a cross section of a TSN and an inclusion containing an inclusion. **E)** shows the 3D structure of this region with the villous cytotrophoblast in yellow and the syncytiotrophoblast in green. **F)** shows a higher magnification image of the structure of the villous cytotrophoblast derived inclusion and the TSN channel. BM and MVM are the basal membrane and apical microvillous membranes of the syncytiotrophoblast.

On the basal membrane, continuous and near-continuous TSN were observed opening adjacent to basal lamina in 16 cases and regions adjacent to cytotrophoblast in 14 cases.

Of the ten continuous TSNs, eight contained membrane-bound inclusions which, surrounded by pore membrane, formed double-membrane structures (figure 4). Inclusions were also observed in the majority of near-continuous and unilateral nanopores. In three cases, the inclusions appeared to be trophoblastic in origin. In the first case, the inclusion on the apical side appeared to be connected to the syncytiotrophoblast by at least one thin stalk of cytoplasmic material (figure 4a-c). At the available resolution, it was not clear whether other inclusions were also engulfed syncytiotrophoblast. In two cases, an inclusion on the basal side of the placenta was a clear-cut protrusion from an underlying cytotrophoblast cell (figure 4d-f).

Another less commonly observed feature associated with TSNs were thin nanopores within the syncytiotrophoblast, where two closely adjacent membranes were joined by desmosome like adhesions (figure 5). These were observed in two TEM images and one SBFSEM image stack. In one case a desmosome associated nanopore was seen almost crossing the placenta from the basal membrane to near the microvillous membrane (figure 5a). Desmosome associated nanopores were observed in one SBF SEM image stack where they were found to be ribbon or sheet-like structures 5-16 nm wide and estimated at 200-800 nm deep (figure 5b). Desmosome containing nanopores could be seen appearing and disappearing within the same 2D section, which is consistent with a pore rather than a cell-cell junction. Topologically a cell-cell junction would need to either interact with two cell or image boundaries or form a circular feature (figure 5c).

**Figure 5.**
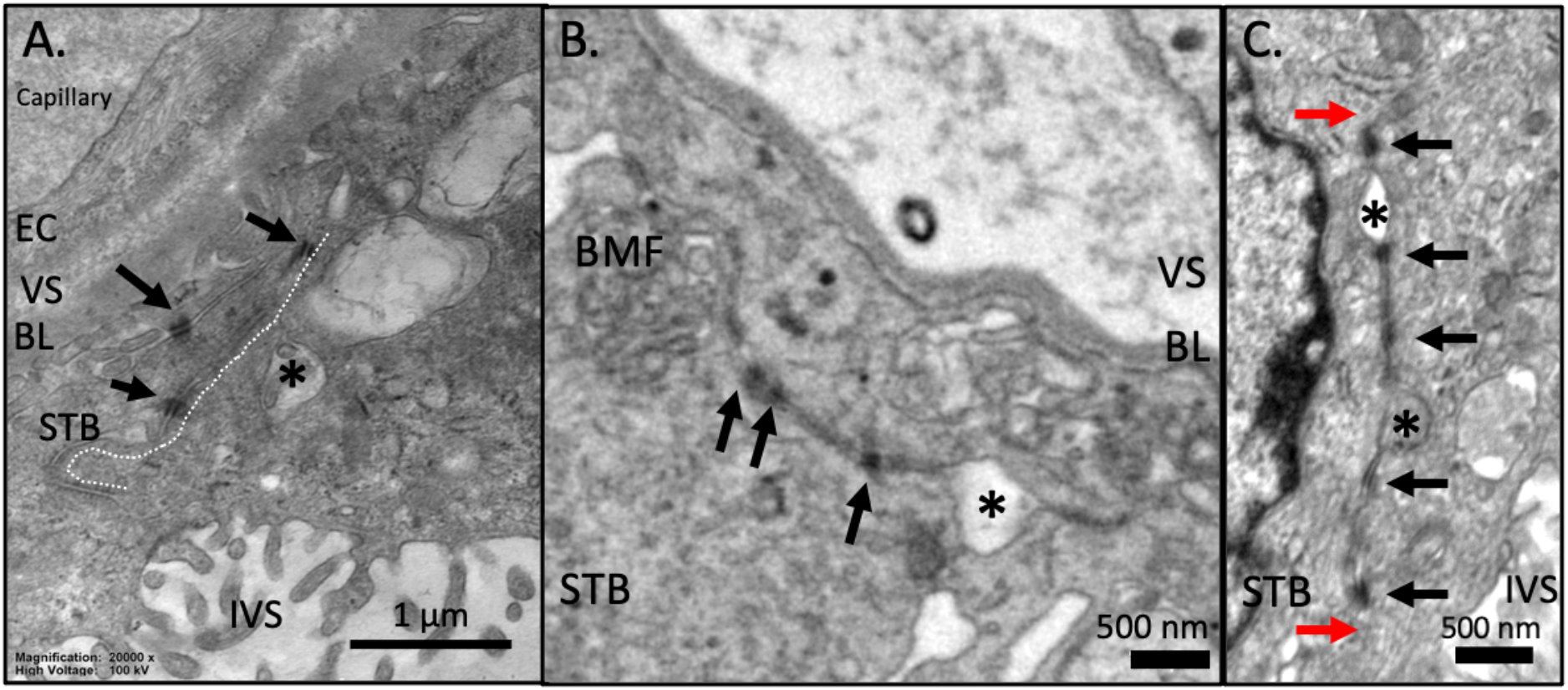
Desmosome associated nanopores. **A)** a TEM image of a desmosome associated nanopore traversing most of the width of the syncytiotrophoblast (white dotted line). The desmosomes are indicated by black arrows. This type of nanopore was associated with dilations typical of the TSNs described in the previous figures (indicated by *). **B)** an SBF SEM image of a desmosome associated nanopore connecting basal membrane folds to a TSN like opening (indicated by *). **C)** A TEM image of a desmosome associated nanopore which begins and ends in the same field of view (red arrows). This demonstrates that these structures are not a cell-cell junction, as if this were the case it would need to intersect 2 sides of the image or be a circle. This channel contains an empty TSN like dilation (indicated by the top*) and one which contains inclusion material (indicated by the bottom*). EC = endothelial cell, VS = villous stroma, BL = basal lamina, STB = syncytiotrophoblast, BMF = basal membrane folds. IVS = intervillous space.

### Modelling diffusion through TSNs

Depending on their geometry, individual nanopores display a large variation in estimated effective *A_pore_/L_pore_* from 0.14 × 10^−8^ m to 2.08 × 10 ^−8^ m (figure 2k). The effective *A_pore_/L_pore_* ratio based on the results of the computational simulations of the ten pores, as depicted in figure 2, was in the order of 10 nm (8.5 ± 7.2 × 10^−9^ m, mean ± SD). By multiplying the mean effective *A_pore_/L_pore_* ratio with the diffusion coefficient for particular solutes using Eq. 2, the average permeability surface area products for a single pore could then be calculated (table 1). Dividing the experimentally observed placental permeability surface area products (from the literature) by the average permeability surface area product for a single pore, resulted in placental pore number estimates between 13 and 37 million pores per gram.

**Table 1:** Estimated number of pores per gram required to explain the experimentally observed placental permeability surface area products.

| | MW<br>(da) | $D$<br>($\text{m}^2 \text{s}^{-1}$) | $PS_{\text{pore}}$<br>($\text{m}^3 \text{s}^{-1}$) | $PS_{\text{placenta}}$<br>($\text{m}^3 \text{s}^{-1} \text{g}^{-1}$) | number of<br>pores ($\text{g}^{-1}$) |
| --- | --- | --- | --- | --- | --- |
| Inulin* | 6179 | $2.6 \times 10^{-10}$ | $2.20 \times 10^{-18}$ | $0.28 \times 10^{-10}$ | $13 \times 10^6$ |
| CrEDTA* | 344 | $7.0 \times 10^{-10}$ | $5.92 \times 10^{-18}$ | $1.06 \times 10^{-10}$ | $18 \times 10^6$ |
| Mannitol* | 182 | $9.9 \times 10^{-10}$ | $8.37 \times 10^{-18}$ | $2.50 \times 10^{-10}$ | $30 \times 10^6$ |
| Creatinine** | 113 | $12.9 \times 10^{-10}$ | $10.9 \times 10^{-18}$ | $4.00 \times 10^{-10}$ | $37 \times 10^6$ |

\*Placental permeability surface area product data taken from Bain *et al.* (1990) [6] and diffusion coefficients from Atkinson *et al.* (1991) [28]. \*\* Placental permeability surface area product based on the average of values from Cleal *et al.* (2007)[29], Edwards *et al.* 1993 [30] & Brownbill *et al.* 1995 [31] and diffusion coefficient from Collins *et al.* (1979) [32].

## Discussion

The demonstration of nanoscale pores punctuating the syncytiotrophoblast challenges our understanding of the placenta as a barrier. It suggests that rather than being a continuous physical barrier between mother and fetus, the syncytiotrophoblast is a molecular sieve facilitating diffusion of small solutes via active maintenance of TSNs. Computational modelling diffusion though these channels suggests that these solutes may include nutrients and toxins, so this transfer route may have profound implications for the fetus.

TSNs were structurally heterogeneous, including simple tubes, tubes with dilated regions and branched structures with blind ends. Many TSNs also contained inclusions of membrane-bound material, in three cases trophoblastic in origin but in most cases the origin was unclear. Some TSNs had multiple openings to the syncytiotrophoblast apical microvillous or basal membranes. The near-continuous and unilateral nanopores are likely in the process of formation, degradation or remodelling. It is also possible that near-continuous TSNs are continuous but that the connections were too thin to observe. The relatively large number of near-continuous and unilateral nanopores is consistent with TSNs being dynamically remodelled.

There was a wide range in calculated permeability parameters for the modelled TSNs, with diffusion through the most permeable being 10 × greater than the least permeable. As might be expected, shorter, wider TSNs had the greatest permeability while the longer TSNs had the lowest permeability (excluding the TSN with multiple apical and basal openings).

The time scale for diffusion 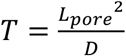 through the TSNs is < 60 ms for the compounds studied. As such TSNs will operate under quasi steady state conditions, given that the timescale of diffusion through the TSN is very fast compared to any changes in concentration in the much larger volumes of the maternal intervillous space and fetal capillary to either side.

There are differences in the placental permeability of different compounds from literature, which could not be fully explained based on diffusivity and thus affected the number of pores estimated, with higher estimated pore numbers for smaller molecules. One explanation for this is that there may be more thinner pores available that only allow passage of smaller molecules. However, we do not think that this is the explanation for what is observed here. Paracellular diffusion across the placenta is size-selective, but the TSNs are likely to be too wide to impose this size selectivity on smaller molecules. Physiological estimates of placental pore sizes in rodents are in the order of 17 nm, which is consistent with the thinnest regions of the nanopores described here and with the desmosome associated channels we observed [15]. For comparison glucose is 1.5 nm long so would be expected to diffuse freely though pores of this size. If the TSNs do not impose size selectivity, other structures such as the trophoblast basal lamina or the capillary endothelium might, and further research is required to understand this better [16].

Molecules such as IgG or nanoparticles such as ultrafine diesel exhaust could diffuse into most regions of the TSNs, but the diameter would sterically hinder their transfer. TSN diameter might be a significant barrier to the transfer of large molecules unless an additional biological mechanism facilitates this transfer.

On the basal membrane TSNs opened adjacent to both basal lamina and cytotrophoblast cells. TSNs opening to the basal lamina would provide the most direct pathway to the fetal circulation. Substances diffusing through TSNs opening adjacent to cytotrophoblast would have a less direct path to reach the fetus but these solutes could interact with the cytotrophoblast themselves.

Further research is required to determine TSN densities in placental tissue. An initial approximation for the TSN density was obtained by dividing the number of observed TSNs (10 continuous or 30 including continuous and near-continuous TSNs) by the imaged volume of 0.00000017 cm^3^. This initial approximation for TSN density is 58 million TSNs per cm^3^, and including near-continuous TSNs is 162 million/cm^3^. This estimate is in line with modelled estimate of the number of TSNs required to mediate transfer of known paracellular markers such as inulin and creatinine.

The 50 nm z-resolution is a potential technical limitation in this study. Firstly, it may lead us to classify some continuous TSNs as near continuous and secondly could also affect the accuracy of the geometric representation in the computational models affecting the calculated permeability parameters. The use of focused ion beam scanning electron microscopy could reduce z depth, but this approach has a smaller field of view making finding TSNs more difficult.

The physiological role of TSNs is likely to include the maintenance of ionic and osmotic homeostasis between mother and fetus. Diffusion of ions and bulk fluid flow would balance osmotic and pressure gradients that could build up across an impermeable placenta. TSNs may also facilitate the placental transfer of nutrients, such as glucose, with maternal to fetal gradients to the fetus [17]. However, the physiological roles of TSNs may come at the cost of allowing non-selective transfer of potentially harmful substances.

The presence of membrane-bound inclusions within the TSNs suggests their role could be more complex than simply being pores for diffusion. The nanopore inclusions and surrounding TSN membrane create a double membrane structure which show ultrastructural similarities to autophagosomes [18]. Nanopore inclusions can be observed in a previous study that sought to identify TSNs using lanthanum perfusion [7].

While the origin and nature of the inclusions remains unclear, in some cases they appeared trophoblastic in origin. One inclusion appeared to be connected to the syncytiotrophoblast by a stalk of cytoplasmic material which is consistent with syncytiotrophoblast being pinched off into a nanopore to form an inclusion. If this interpretation is correct this could potentially represent the initiation of autophagy or material that will be shed from the placenta. In other cases, inclusions on the basal side of the placenta were observed to be protrusions from underlying cytotrophoblast cells.

A rare feature associated with TSNs were desmosome associated nanopores which could be observed arising from basal membrane folds [19], and within the cytoplasm but not observed connecting to the microvillous membrane. The connect to dilations that appear similar to, or are, the TSNs that are the main focus of this paper. It is possible that the TSNs represent formation or breakdown of TSNs or that they are a separate class of nanopore that intersect with TSNs. The desmosome associated nanopores have been observed previously in 2D TEM images where it was suggested that they are remnants of cytotrophoblast-syncytiotrophoblast fusion. We cannot exclude this trophoblast fusion explanation, but in the limited number of 3D examples we identified the structures were more like ribbon than a cell-cell junctions [20]. Desmosome associated nanopores are very thin and so may not be visible when the block is in the wrong orientation and so could be more common than they appear.

Cells that cross or penetrate epithelial barriers have been observed to express tight junction proteins to facilitate movement through cell-cell junctions, and the desmosome containing nanopores in the syncytiotrophoblast could provide a route and a mechanism by which maternal or fetal cells could cross the syncytiotrophoblast [21, 22]. Images of erythrocytes protruding through the syncytiotrophoblast are consistent with distensible channels that could allow cell transfer [23].

There is considerable diversity in placental structures across species and given the possibility that the syncytiotrophoblast has evolved independently in different branches of the evolutionary tree, the diversity of TSNs (or similar structures) is of interest. To date, full width trans-syncytial channels have only been described in the degu which has both short direct channels in thin regions of syncytiotrophoblast and more complex channels connecting infoldings of basal and apical surfaces [24]. The human TSNs had a distinct appearance but were most similar to the more complex Degu channels, although longer and thinner. The distribution of TSNs across species may inform their biological role and origins.

Identifying the molecular processes underlying TSN formation will be key to understanding these structures. While TSNs may form by a unique mechanism, they are more likely to co-opt at least some of the known molecular mechanisms that mediate endocytosis. There are multiple endocytic processes, including clathrin and caveolin, micropinosomes or CLIC/GEEC, which form tubular structures [25]. The CLIC/GEEC may be of particular interest as these form tube-like structures [26].

In conclusion, this study has demonstrated the existence of TSNs providing a non-selective diffusion pathway across the placenta. These TSNs may allow pharmacological drugs, environmental toxins and even particulate pollutants across the syncytiotrophoblast. Accurately determining the density of TSNs in healthy pregnancy is necessary to confirm the capacity of TSNs to mediate transfer. Furthermore, establishing the density of TSNs in disease may provide novel insights into disease processes.

## Acknowledgements

This work was supported by BBSRC project grant BB/R002762/1. Equipment in the Biomedical Imaging Unit was supported by MR/L012626/1 Southampton Imaging under MRC UKRMP Funding.

## Competing interests

The authors have no competing interests to declare

## Notes

https://dx.doi.org/10.6019/EMPIAR-10967

### Competing Interest Statement

The authors have declared no competing interest.

### Summary of Updates

Revisions to address reviewer feedback.

